# CryoSTAR: Leveraging Structural Prior and Constraints for Cryo-EM Heterogeneous Reconstruction

**DOI:** 10.1101/2023.10.31.564872

**Authors:** Yilai Li, Yi Zhou, Jing Yuan, Fei Ye, Quanquan Gu

## Abstract

Resolving conformational heterogeneity in cryo-electron microscopy (cryo-EM) datasets remains a significant challenge in structural biology. Previous methods have often been restricted to working exclusively on volumetric densities, neglecting the potential of incorporating any pre-existing structural knowledge as prior or constraints. In this paper, we present a novel methodology, cryoSTAR, that harnesses atomic model information as structural regularization to elucidate such heterogeneity. Our method uniquely outputs both coarse-grained models and density maps, showcasing the molecular conformational changes at different levels. Validated against four diverse experimental datasets, spanning large complexes, a membrane protein, and a small single-chain protein, our results consistently demonstrate an efficient and effective solution to conformational heterogeneity with minimal human bias. By integrating atomic model insights with cryo-EM data, cryoSTAR represents a meaningful step forward, paving the way for a deeper understanding of dynamic biological processes.^1^

## Introduction

Single particle cryo-electron microscopy (cryo-EM) is a structural biology tool that can directly observe the conformational heterogeneity of each biomolecule, that each dataset contains many 2D projections of 3D structures from potentially different conformational states^1^. Traditional algorithms (e.g., 3D classification) treat the heterogeneity in the dataset as discrete clusters and assign each particle to the best class^2–6^. However, in many real datasets, heterogeneity often comes from conformational dynamics, a continuous process. Using traditional algorithms often results in the 3D density maps blurry in the flexible regions.

A few algorithms in recent years have been developed to resolve continuous heterogeneity from cryo-EM datasets. For example, principal component analysis (PCA) and its variants have been used to describe the variability within the dataset, which model the heterogeneity as a linear combination of a few bases^7–10^. To achieve more expressive power with nonlinearity, deep learning-based methods were developed to map such heterogeneity onto nonlinear manifold embeddings. For example, cryoDRGN^11^ and cryoDRGN2^12^ use a variational autoencoder (VAE)^13^ based approach to map the heterogeneity within the dataset to a latent space. A generative decoder is used to generate a 3D volume given a sampled point from the latent space. On the other hand, 3DFlex^14^ explicitly models the motion of flexible regions by learning a 3D deformation field and optimizing a canonical density, while encouraging local smoothness and rigidity. Nevertheless, these methods approach the continuous heterogeneity issue solely from a computer vision perspective, without leveraging any prior knowledge that could be used as structural constraints.

Some recent works tried to incorporate information from the atomic model into the pipeline, or to directly output coarse-grained (CG) atomic models for better interpretation. For example, Chen et al.^15^ explore the possibility of explicitly modeling atoms or residues with a Gaussian density as a follow up work of *e2gmm*^16^. Other methods try to decompose heterogeneity into a few bases using normal mode analysis (NMA)^17,18^ or Zernike polynomials^19^. The atomic-level or residue-level information often helps provide interpretation by offering sensible models with motions. However, these methods either only find relatively small continuous motions^17,19^, or are only verified on synthetic data^18,20,21^. In particular, NMA is more suitable for finding the possible fluctuation rather than representing the real, complex motion^22^.

In this paper, we introduce cryoSTAR (**<u>St</u>**ructur**<u>a</u>**l **<u>R</u>**egularization), a deep neural network model that resolves continuous conformational heterogeneity from cryo-EM datasets by generating both density maps and reasonable coarse-grained (CG) models for different conformations. Our method requires an initial atomic model as the reference, whose structural information is used to properly regularize the inferred conformational dynamics. This enables us to correctly preserve the local structures, narrowing the search space by avoiding fallacious solutions, achieving better and faster convergence. The learned conformational heterogeneity allows for the concurrent generation of density maps and coarse-grained models. Notably, the density map can be used to evaluate and validate the conclusions from the coarse-grained models.

We validate cryoSTAR on a synthetic dataset with known ground truth (Methods and Extended Data Fig. 1) and apply it to four public experimental datasets. On the pre-catalytic spliceosome^23^ (EMPIAR-10180) and the U4/U6.U5 tri-snRNP^24^ (EMPIAR-10073), we recover the motions that generally agree with other methods^11,14^, and generate both density maps and reasonable coarse-grained models with the corresponding dynamics. We further demonstrate the effect of using incomplete and slightly erroneous atomic models as reference. Despite the potential biases in the generated coarse-grained model, the density maps are robust and can serve as a reliable tool for validating the coarse-grained models. We then show the effectiveness of cryoSTAR on the TRPV1 channel^25^ (EMPIAR-10059), a small membrane protein. Without excluding the lipid nanodisc region by manual masking or particle subtraction, cryoSTAR successfully resolves its conformational heterogeneity. Finally, we show the performance of cryoSTAR on α-latrocrustatoxin (α-LCT)^26^ (EMPIAR-10827), a small protein with a molecular weight of 130 kDa. We resolve the continuous motion that agrees with the original paper found by discrete 3D classification^26^, and also uncover a different type of flexible motion. Structural regularization is especially beneficial in challenging cases, such as resolving continuous motions in membrane proteins or in smaller proteins. Our experiments suggest that cryoSTAR is a powerful tool for solving the continuous conformational heterogeneity from cryo-EM images.

## Results

### CryoSTAR

CryoSTAR models the conformation heterogeneity in cryo-EM particles as the deformations of a reference atomic model *S*_ref_ ∈ ℝ^*N*×3^, where *N* is the number of residues, and applies proper regularization to ensure the integrity of the structures (Fig. 1a). With a variational autoencoder (VAE)^13^ as the neural network architecture, cryoSTAR compresses the conformational heterogeneity into a latent variable, and deforms a coarse-grained model *S*_ref_, which is pre-computed from a reference atomic model provided as an input (Fig. 1b). Specifically, given an image *I* ∈ ℝ^*D*×*D*^ from the cryo-EM dataset, cryoSTAR uses a VAE to predict the corresponding deformation Δ^*V*^^ that modifies the reference structure to the deformed structure ^*V*^^ = *V*_ref_ + Δ^*V*^^. The deformed model is then converted to a Gaussian density *V* ∈ ℝ^*D*×*D*×*D*^, which is a combination of Gaussian functions mapping the volumetric density map to the CG atomic model (see Methods for details). The 2D projection of this Gaussian density can be computed with the given orientation angles and CTF parameters (Fig. 1b). The projection is compared with the input particle image, regularized by the structural constraints derived from the given atomic model (Fig. 1a).

**Figure 1.**
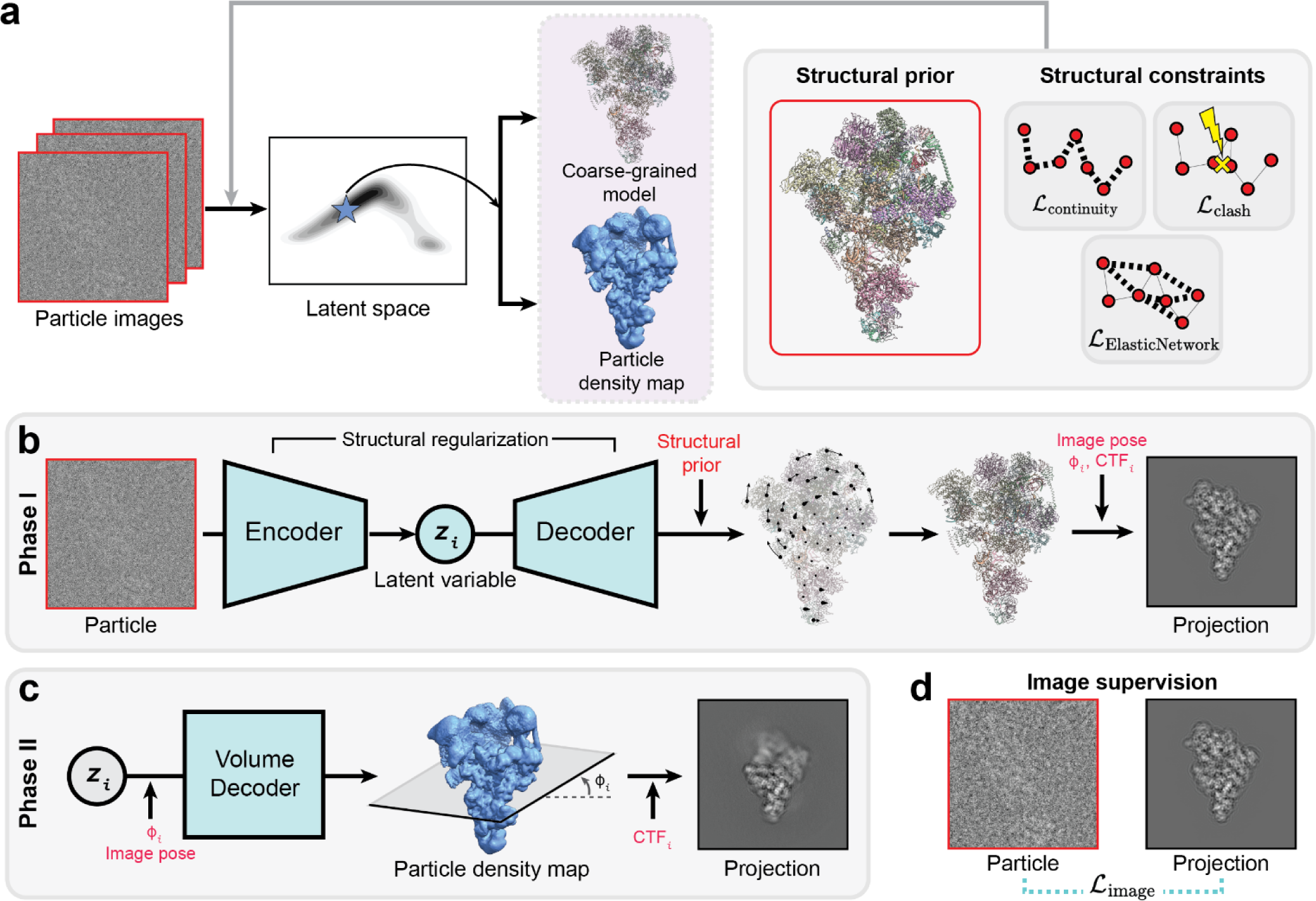
CryoSTAR optimize conformational heterogeneity with structural prior and constraints, and concurrently output both coarse-grained models and density maps with different conformations, **a,** From particle images, cryoSTAR uses a referece atomic model as the structural prior and proper structural constraints to map conformational heterogeneity into a latent space. Both coarse-grained models and density maps of different conformations can be generated concurrently by sampling from the learned latent space, **b,** The training step of cryoSTAR has two phases. In Phase I, a particle image is fed into a variational autoencoder (VAE), where conformational heterogeneity is compressed into the latent variable z. Based on the output of the decoder, the coarse-grained model derived from a reference atomic model will be deformed to the predicted conformation. With the given particle pose and CTF, the predicted projection of the deformed model can be calculated and compared with the input particle. The optimization is under the structural regularization in **a. c,** After the VAE in Phase I **is** fully trained, the latent variable is detached and used to train a volume decoder to obtain a neural network representation of the density maps. Images and volumes are displayed in real space for better visual understanding, d, Optimization in both phase I and II are under the supervision of the input particle. Neural networks and latent variables are colored by light aqua when they are trainable.

### Structural regularization in cryoSTAR

CryoSTAR requires a reference atomic model as its structural prior, which differs from most existing methods. Incorporating the atomic model into heterogenous reconstruction allows imposing meaningful constraints on the potential motions of the target. This structural prior and the constraints are pivotal for tackling the conformational heterogeneity and distinguish cryoSTAR from other methods. In particular, the atomic model assists the algorithm in filtering out evidently incorrect dynamics, thereby facilitating better solutions and rapid convergence.

CryoSTAR uses a structural regularization under three basic assumptions (Fig. 1a):

1. Two adjacent residues in the same chain should always remain connected. CryoSTAR uses a continuous loss ℒ_cont_ to enforce this:

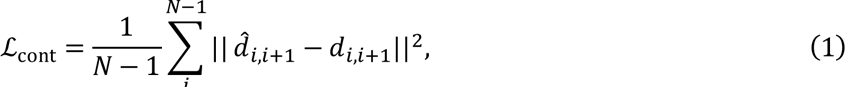

where *d*_*ij*_ and ^*d*^^_*ij*_ denote the distance between the *i*-th and *j*-th residues in the reference structure *S*_ref_ and the predicted structure ^*V*^^, respectively. 1. 2. Residues should not become too close after the predicted deformation. CryoSTAR uses a clash loss ℒ_clash_ to penalize if clashing happens:

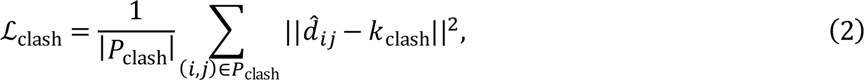

where *P*_clash_ denotes the set of residue pairs that experience collision during training, (*i*, *j*) is an index pair numbering the residues in the structure, and the constant value *k*_clash_ is a predetermined cutoff.

1. 3. Local structures should be as rigid as possible. CryoSTAR builds an elastic network (EN) from the reference atomic model, and uses an elastic network loss ℒ_EN_ to encourage this local rigidity:

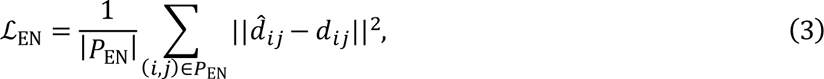

where *P*_EN_ is a set of edges for building the elastic network. Since the elastic network may be subject to changes during conformational dynamics, cryoSTAR adaptively selects the edges presented in the elastic network for regularization in training (see Methods for details).

To summarize, the structural regularization in cryoSTAR enforces the continuity of a chain, prevents the residues from clashing, and encourages local rigidity with an adaptive elastic network (Fig. 1a). This structural regularization is critical for the model to output reasonable coarse-grained models and recover the correct dynamics.

### Generating density maps with minimal bias

After the VAE is fully trained in Phase I, the latent variable is extracted and used to train a volume decoder to obtain a neural network representation of the densities in Phase II (Fig. 1c). The optimization of the density maps is solely guided by the input particles (Fig. 1d), remaining unaffected by the structural prior and regularization. This minimizes the reference bias on the output density. Therefore, the generated density maps can be used to evaluate and corroborate the coarse-grained models from Phase I. As a result, cryoSTAR can simultaneously generate both reasonable coarse-grained models and volumetric density maps at different conformations (Fig. 1a), helping the users evaluate and interpret the results at different levels.

### CryoSTAR finds conformational heterogeneity of large complexes

The yeast pre-catalytic B complex spliceosome is a big complex with more than 10,000 residues including amino acids and nucleotides. The cryo-EM dataset (EMPIAR-10180) includes conformational dynamics on the Sf3b and the helicase regions (Fig. 2a), as resolved by other methods^11,19^, and has been used as a benchmark to test continuous heterogeneity algorithms. Such continuous motions are impossible to uncover by traditional 3D classification, which classifies the particles into discrete clusters of conformation. We use the atomic model in Plaschka et al.^23^ (PDB: 5NRL) as the reference atomic model to train cryoSTAR. By traversing along the first principal component of the latent space (Fig. 2b), cryoSTAR reveals the motion of the SF3b and the helicase regions (Fig. 2a, 2c and Supplementary Video 1). While the SF3b can bend towards the “body”, the helicase region can curve down to the “foot”. This result is generally consistent with the results found by other methods^11,19^. Furthermore, the generated coarse-grained models do not exhibit unnatural deformations or disruptions, and more importantly, their motion is corroborated by the particle density maps (Fig. 2a and 2c). Compared to the other method that also outputs atomic models^19^, our result shows a more obvious motion and more reasonable coarse-grained models without local shearing effects, e.g., the local structures of the alpha helices remain intact. For example, our findings suggest that the alpha helix of Spp381 shifts toward the foot domain, which can be verified by the associated density maps, showing a seamless transition without any discernible artifacts (Fig. 2c). The reference atomic model (PDB: 5NRL) also contains the U2 snRNP region, which exhibits weaker density in the consensus map, potentially due to compositional heterogeneity^11^. This region is also evident in the particle density maps produced by cryoSTAR, with a similar diminished intensity compared to other areas. CryoSTAR suggests a possible motion of the U2 core in the generated coarse-grained models; however, such movement cannot be fully supported by the particle density maps, which do not reveal obvious motions (Extended Data Fig. 2). Resolving the dynamics at different levels provides the users with rich information for better evaluation and interpretation.

**Figure 2.**
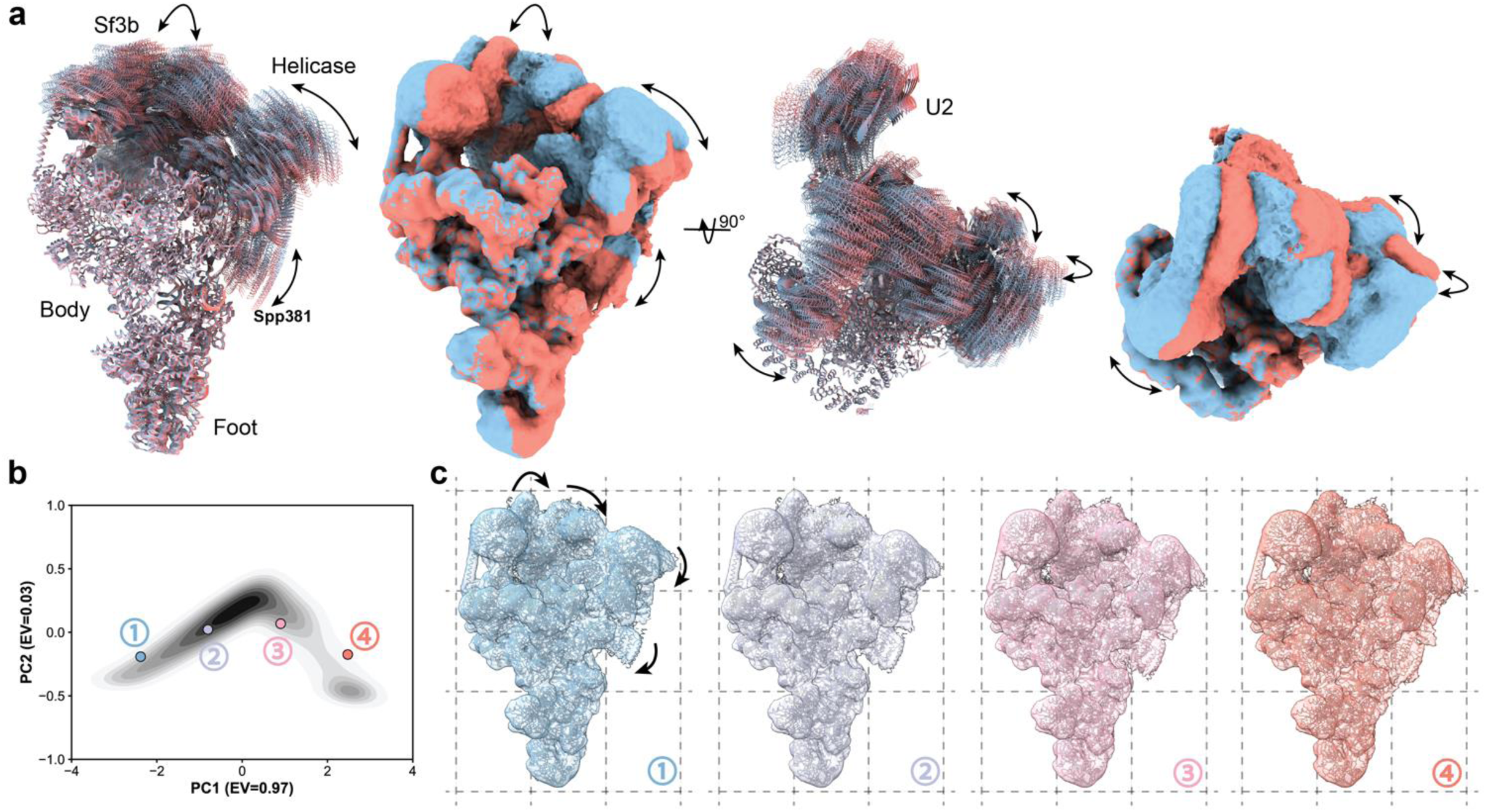
CryoSTAR reveals the motion of Sf3b and helicase domains in the pre-catalytic spliceosome dataset (EMPIAR-10180). **a,** Colored series of ten coarse-grained models and two particle density maps generated by cryoSTAR, sampling along the first principal component of the latent space, **b,** PCA visualization of the cryoSTAR latent space, where the color depth represents the particle population, **c,** Four coarse-grained models and particle density maps generated by sampling along the first principal component in the latent space, as marked in **b** with the corresponding colors. All density maps are shown using the same isosurface levels.

The U4/U6.U5 tri-snRNP is a considerable part of the spliceosome before activation and is known to have flexible regions especially in the head and arm parts^14,24^ (Fig. 3a), which cannot be resolved by traditional 3D classification. An essentially complete atomic model (PDB: 5GAN) was previously built from the consensus density map solved by cryo-EM^24^. Using this atomic model as the input, we train cryoSTAR on the 138,899 particles in the EMPIAR-10073 dataset. We then sample along the first and second principal component of the learned latent space (Fig. 3b and Extended Data Fig. 3b), from which we generate the corresponding coarse-grained models and density maps (Fig. 3c and Extended Data Fig. 3c). Notably, cryoSTAR resolves the motion of the head domains of tri-snRNP in this dataset, which can bend towards the foot domain, as supported by the coarse-grained models and density maps (Fig. 3c, Extended Data Fig. 3c and Supplementary Video 2). Moreover, cryoSTAR also finds a possible rotation of the arm domain along the first principal component of the latent variable, which is uncorrelated with the bending of the head domain (Extended Data Fig. 3c and Supplementary Video 2). Nevertheless, the generated density maps do not exhibit a sufficiently close alignment to substantiate this finding, necessitating further investigation.

**Figure 3.**
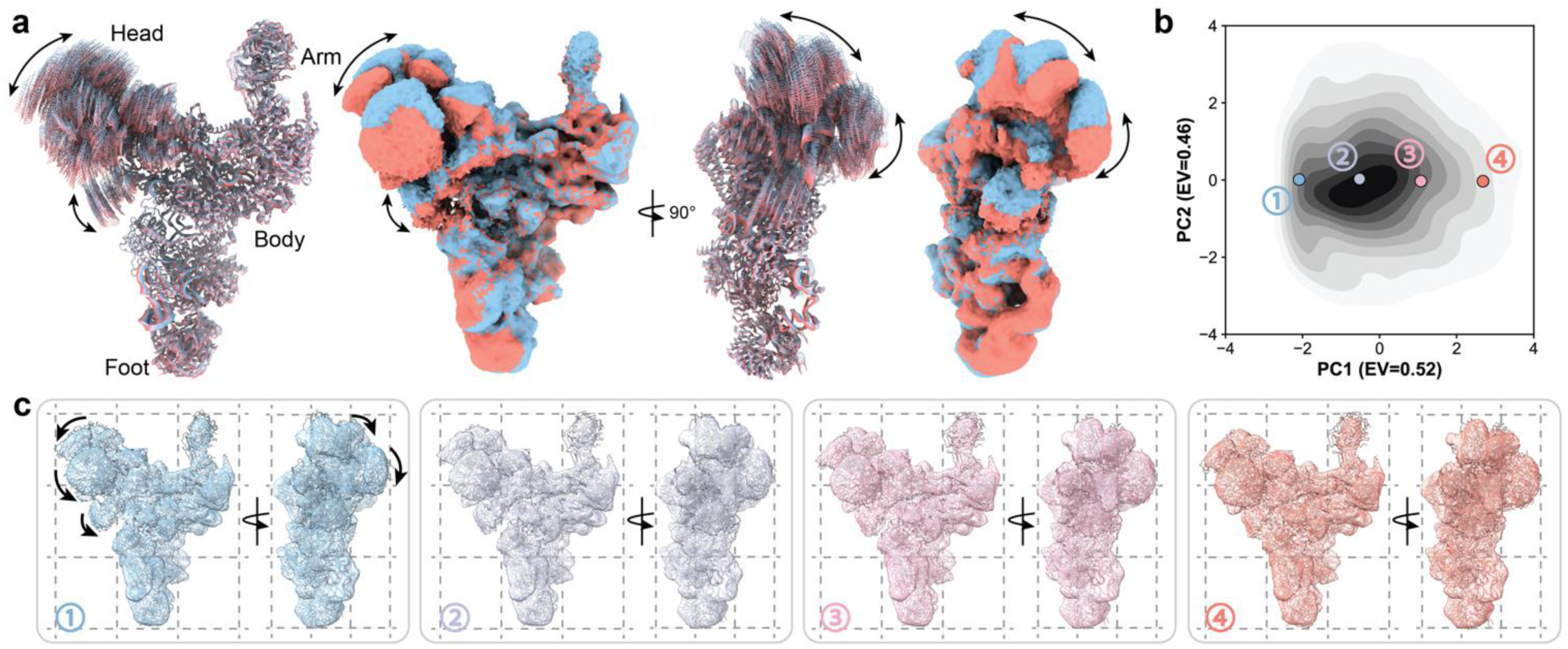
CryoSTAR uncovers the motion of head domain of the U4/U6.U5 tri-snRNP dataset (EMPIAR-10073). **a,** Colored series of ten coarse-grained models and two particle density maps generated by cryoSTAR, sampling along the first principal component of the latent space, **b,** PCA visualization of the cryoSTAR latent space, where the color depth represents the particle population, **c,** Four coarse-grained models and particle density maps generated by sampling along the first principal component in the latent space, as marked in **b** with the corresponding colors. All density maps are shown using the same isosurface levels.

### CryoSTAR is robust to imperfect reference atomic models

Using an atomic model as the initial reference may incorporate a strong bias in data processing. To test the robustness of cryoSTAR to imperfect reference atomic models, we manually edit the input atomic models of both the yeast pre-catalytic spliceosome and the U4/U6.U5 tri-snRNP and evaluate if the results are consistent. For the pre-catalytic spliceosome, we manually remove SmG in U5 snRNP (71 amino acids) from the input atomic model (Fig. 4a), and use the edited model as the input reference atomic model to train cryoSTAR. We find the resolved dynamics match those using the complete atomic model as the reference (Fig. 4b). The generated coarse-grained models show a slight bias that the neighboring amino acids are shifted to fill in the density with the missing residues. Nevertheless, the density maps are consistent with those using the complete atomic model, even in the regions where the corresponding residues are missing (Fig. 4c). Similarly, we manually rotate Prp31 (415 amino acids) in the tri-snRNP atomic model by 45 degrees (Fig. 4d) and use the edited model as the reference to train cryoSTAR. The uncovered motions and the density maps are consistent with the results using the unedited model (Fig. 4e and 4f). Moreover, we find that Prp31 in the generated coarse-grained models overcomes the bias from the reference atomic model and positions itself accurately (Fig. 4f). This suggests that cryoSTAR is not only robust to imperfections in reference atomic models but can also rectify them in some circumstances. However, it should be noted that cryoSTAR needs the reference atomic model to cover the flexible regions, without which the dynamics cannot be correctly modeled.

**Figure 4.**
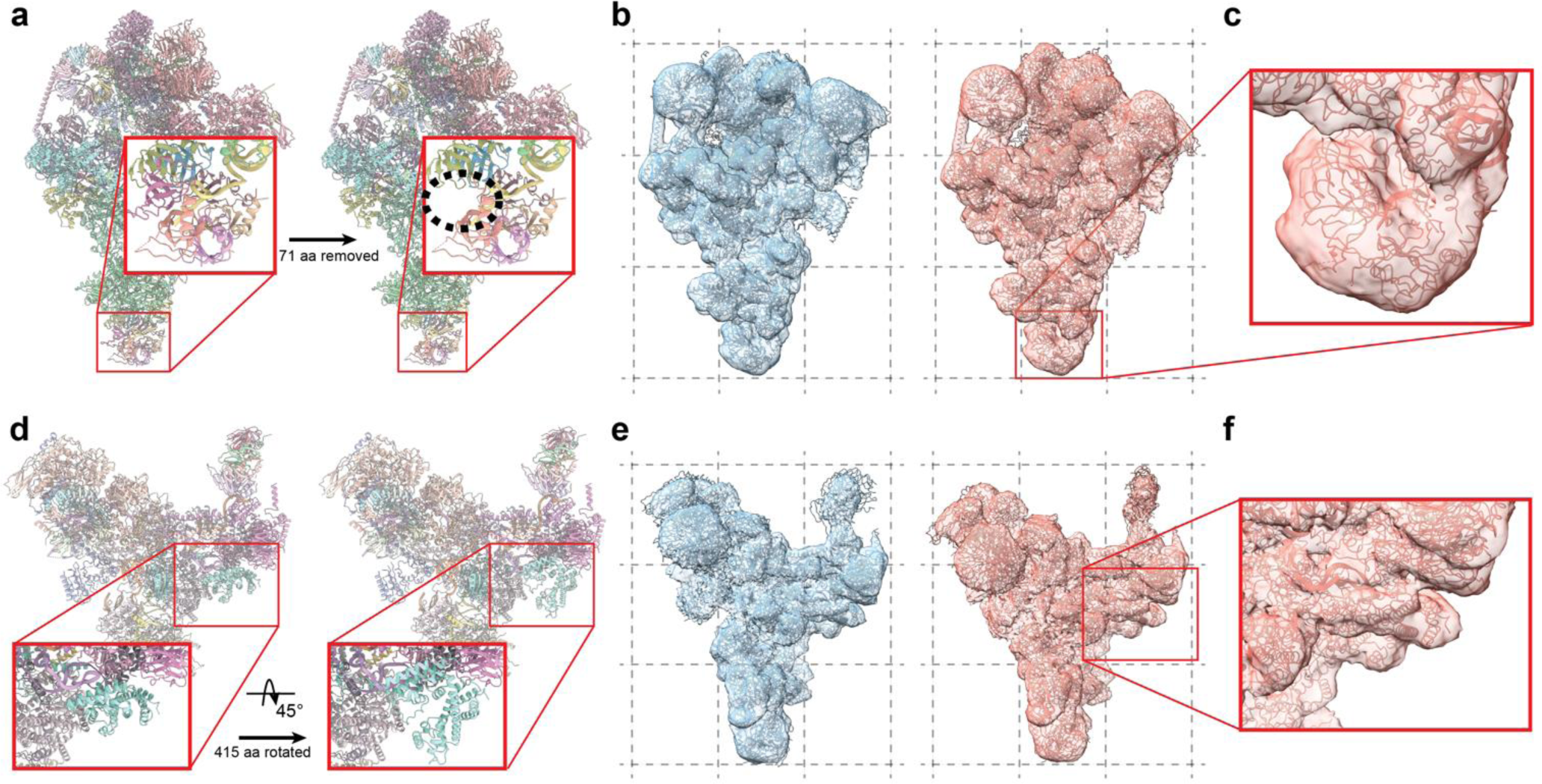
Density maps generated by cryoSTAR are robust to bias from imperfect reference atomic models, **a,** The original (PDB: 5NRL) and the edited reference atomic models of the pre-catalytic spliceosome. SmG in U5 snRNP (71 amino acids) was removed from the original model and used as the reference atmoic model for cryoSTAR. **b,** The coarse-grained models and particle density maps generated by sampling at the 5-th and 95-th percentile of first principal component of the latent space, **c,** Right of **b,** zoomed in to show the details of the edited region. While the coarse-grained model tries to fill the missing residues, the particle density map remains unbiased, **d,** The original (PDB: 5GAN) and the edited reference atomic models of the U4/U6.U5 tri-snRNP. Prp31 (415 amino acids) was manually rotated by 45 degrees and used as the reference atmoic model for cryoSTAR. **e,** The coarse-grained models and particle density maps generated by sampling at the 5-th and 95-th percentile of first principal component of the latent space, **e,** Right of **f,** zoomed in to show the details of the edited region. The coarse-grained model overcomes the reference bias and finds the correct position and pose of the edited chain. The generated particle density also supports the coarse-grained model.

### CryoSTAR reveals membrane protein dynamics

We further explore the application of cryoSTAR to a smaller membrane protein. TRPV1 functions as a heat sensory ion channel and is a tetrameric membrane protein with a molecular weight of 380 kDa. Starting from the polished particles in EMPIAR-10059^25^, we first obtain the particle poses using homogenous refinement in cryoSPARC^3^ without imposing symmetry. The atomic model built from this dataset (PDB: 5IRX)^25^ is not complete and does not cover the regions that are suspected to be flexible, thus we use another atomic model (PDB: 7RQW)^27^ as the reference to train cryoSTAR. While the TRPV1 particles in this dataset were solubilized in nanodisc, we do not use any separate masks or particle subtraction to remove the nanodisc signal in the particle.

CryoSTAR finds a subtle yet smooth motion of the peripheral soluble domains in the protein, as they can move closer or away from the center of the channel (Fig. 5 and Supplementary Video 3). Since no symmetry has been imposed, the motion of each subunit is resolved independently and can be visualized by traversing along the different principal components (Fig. 5 and Supplementary Video 3). Compared to the generated densities, the coarse-grained models can occasionally exaggerate the motion, leading to misalignment with the density map (Fig. 5e). Nevertheless, since the density maps are derived directly from particle images, they remain unaffected by the coarse-grained models, serving as a reliable benchmark to assess the coarse-grained model results (Fig. 5e and 5f). This can be further supported by the fact that the generated density maps include the densities of the detergent micelle, whereas the coarsegrained models do not (Fig. 5d).

**Figure 5.**
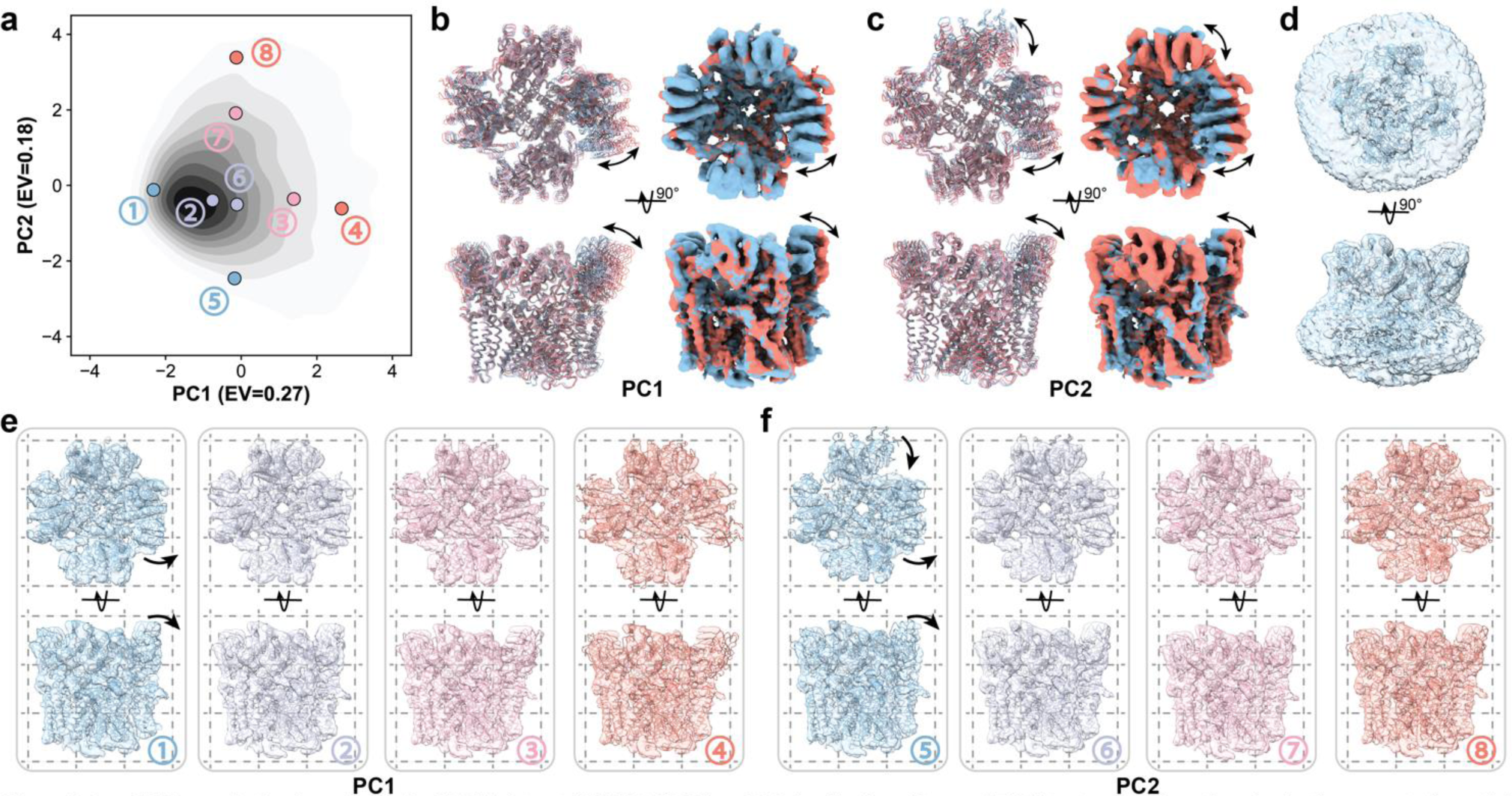
CryoSTAR reveals the dynamics in the TRPV1 dataset (EMPIAR-10059). **a,** PCA visualization of the cryoSTAR latent space, where the color depth represents the particle population, **b-c,** Colored series of ten coarse-grained models and two particle density maps (top and side views) generated by cryoSTAR, sampling along the first and second principal component of the latent space, respectively, **d,** Example of the generated coarse-grained model and particle density map displayed at low isosurface level to reveal the density of the lipid nanodisc, **e-f,** Coarse-grained models and particle density maps generated by sampling along the first and second principal component in the latent space, respectively, as marked in a with the corresponding colors and numbers. Motions of the individual subunits can be seen. All density maps are shown using the same isosurface levels.

Structural regularization plays a pivotal role in revealing the dynamics from this membrane protein. Methods without structural prior, such as cryoDRGN, fall short in revealing the hidden dynamics of membrane proteins (Extended Data Fig. 4a). Without particle subtraction, cryoDRGN tends to focus primarily on regions of highest variability, such as the micelle and nanodisc, rather than the protein itself. In contrast, the structural prior allows cryoSTAR to focus on the protein, even without the need for external masks (Extended Data Fig. 4b). This highlights the superiority of structural regularization in uncovering the intricate dynamics of membrane proteins.

### CryoSTAR uncovers the conformational heterogeneity of α-latrocrustatoxin (α-LCT)

Finally, we demonstrate the ability of cryoSTAR to resolve the continuous heterogeneity on a small protein. α-latrocrustatoxin (α-LCT) is one of the seven black widow spider neurotoxins with a molecular weight of 130 kDa^26^. A recent work found two different conformations of α-LCT via discrete 3D classification, where the curvature of the Ankyrin-like Repeat Domain (ARD) alters, so that the tail of ARD can move towards the head domain. We reanalyze the original dataset (EMPIAR-10827)^26^, curate 230,218 particles after 2D classification and obtain particle poses from homogeneous refinement. After training cryoSTAR, we uncover two types of motions of α-LCT (Fig. 6 and Supplementary Video 4). Along the first principal component, the ARD tail moves aside relative to the head domain (Fig. 6a, 6b and 6d). On the second principal component, the ARD tail bends towards or away from the head domain (Fig. 6a, 6c and 6e), which is consistent with the two conformations found by discrete 3D classification^26^. In both cases, the generated coarse-grained models are reasonable and fit well with the density maps (Fig. 6d and 6e). We would like to underline that the original elastic network built from the reference atomic model connects the ARD tail with the head domain; however, the adaptive relaxation of the elastic network (see Methods for details) helps the model to eliminate this unwanted interaction. This mitigates the bias from the reference atomic model and encourages cryoSTAR to explore more possible conformations.

**Figure 6.**
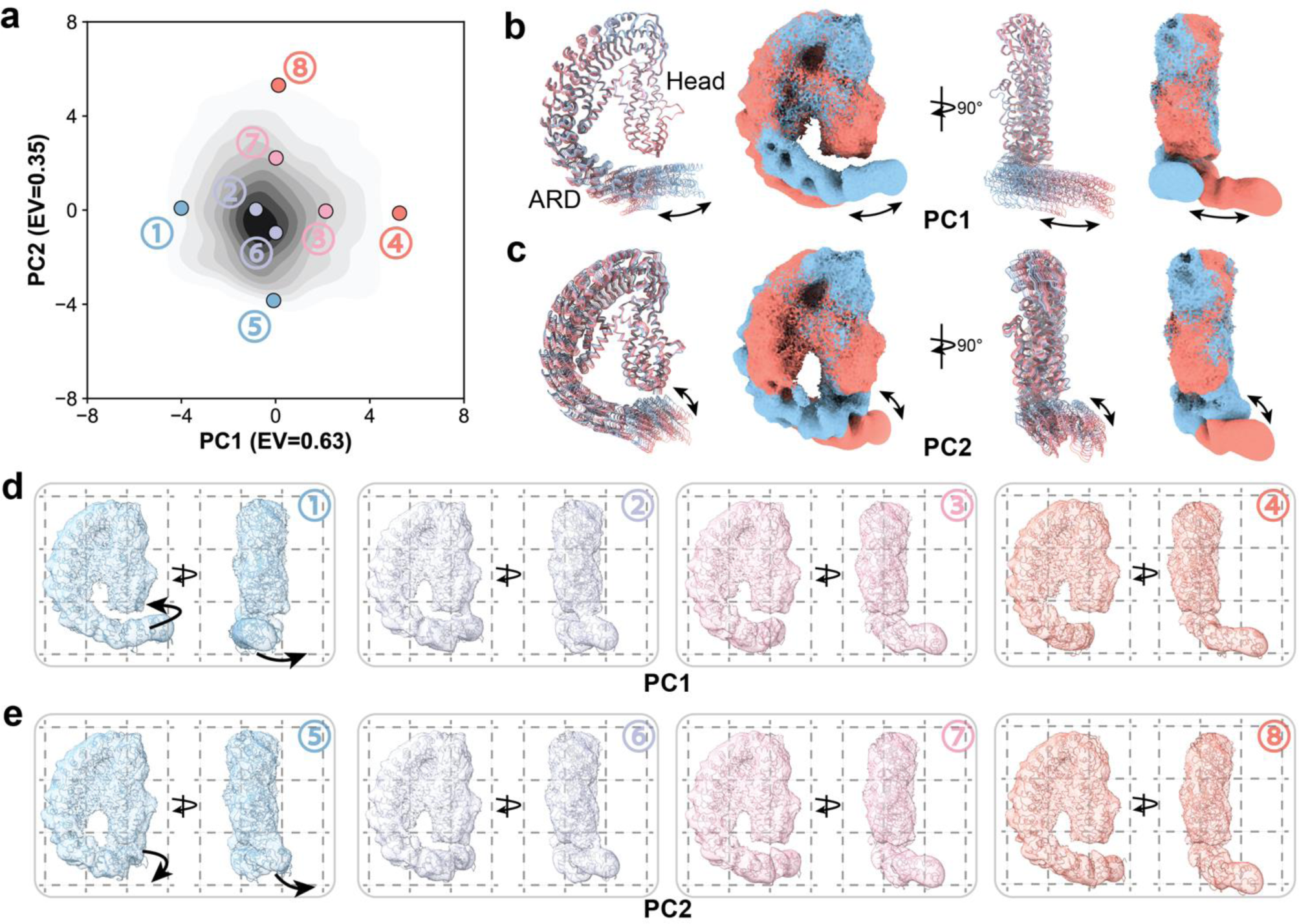
CryoSTAR reveals two different motions of the Ankyrin-like repeat domain (ARD) from the a-LCT dataset (EMPIAR-10827). **a,** PCA visualization of the cryoSTAR latent space, where the color depth represents the particle population, **b-c,** Colored series of ten coarse-grained models and two particle density maps generated by cryoSTAR, sampling along the first and second principal component of the latent space, respectively. **d-e,**The coarse-grained models and particle density maps generated by sampling along the first and second principal component in the latent space, respectively, as marked in a with the corresponding colors. Two different types of motion of the ARD tail are uncovered. All density maps are shown using the same isosurface levels.

The efficacy of structural regularization is particularly evident in this small protein with continuous heterogeneity. For instance, although cryoDRGN can uncover similar dynamic modes, its density maps are less continuous, often missing the ARD tail when traversing the latent space (Extended Data Fig. 5a). In comparison, 3DFlex finds motion that is much less significant, and produces reconstructed density maps that suffer from noticeable artifacts in the ARD tail (Extended Data Fig. 5b). In contrast, the use of structural regularization in cryoSTAR successfully maintains a continuous density throughout movement, effectively circumventing the artifacts in the density maps (Extended Data Fig. 5c).

## Discussion

CryoSTAR offers a distinctive approach in solving conformational heterogeneity from cryo-EM datasets. Uniquely, cryoSTAR resolves such heterogeneity by generating coarse-grained models of different conformations, which are directly learned from the particle images. This innovation allows cryoSTAR to effectively harness the pre-existing structural knowledge from the reference atomic model. This sets our approach apart from other methodologies that typically focus solely on working with volumetric densities. Once the heterogeneity is encoded, the latent embeddings are used to train a volume decoder that can generate density maps of different conformations. These density maps are sourced directly from the particle images and are not influenced by the coarse-grained models. This ensures that they remain relatively unbiased and can be used to cross-reference the structures and dynamics of the coarsegrained models. To sum up, cryoSTAR generates both coarse-grained models and density maps of different conformations, enabling researchers to gain a clearer picture of molecular conformational changes.

It is worth noting that the reference atomic model provided to cryoSTAR serves a crucial role in structural regularization, which limits the searching space to only reasonable conformations during training. This reference model can be derived from fitting or docking in the map, from computational predictions, or any other proper methods. The primary requirement is that it should generally agree with the consensus density. Imperfect reference atomic models have been demonstrated to produce consistent results without significantly biasing the resolved heterogeneity. Importantly, while the coarse-grained models might be subject to reference bias, the generated density maps remain unaffected, serving as a reliable cross-reference. It is also important to note that cryoSTAR does not engage in model building or flexible fitting from a density map with dynamics. Instead, its strength lies in exploiting atomic model information as constraints in solving heterogeneity from the particles. Hence, the generated coarse-grained models lack validation and should be interpreted with caution. Any conclusions drawn from the coarse-grained models should also find support in the corresponding density maps.

Furthermore, it is noteworthy to compare cryoSTAR with methods that derive a canonical density from optimized deformation fields. These methods usually yield density maps with apparent higher local resolution, since the signals from all the particles are contributed to the canonical density ^14^. To prevent overfitting, regularization that encourages local smoothness of the deformation field is usually employed. While being a valid and intuitive constraint, the implementation of “local smoothness” is often arbitrary. This inductive bias is embedded in the deformation field and will subsequently influence the optimization of the canonical density. Consequently, although a reference density map or atomic model is absent, the inductive bias may have a significant impact and cannot be entirely detected with the “gold-standard” Fourier Shell Correlation (FSC) (Extended Data Fig. 5b). In contrast, instead of optimizing one canonical density, the density maps in cryoSTAR are generated independently without the deformation field (Fig. 1c). Therefore, we deliberately minimize this inductive bias, even at the cost of an apparent improvement in local resolution.

While being a powerful method, cryoSTAR has its own limitations. First, cryoSTAR is not designed to solve compositional heterogeneity in the dataset. In such cases, cryoSTAR may generate density maps with weaker densities in certain regions (Extended Data Fig. 2 and Extended Data Fig. 3), but such heterogeneity may not be encoded in the latent embeddings. Second, cryoSTAR requires a reasonably good consensus map so that the particle poses are fairly accurate. For example, although the conformational dynamics of α-LCT can be resolved (Fig. 6), the generated densities are in low resolution and lack structural details, likely due to the inaccurate angular assignments of the particles in the upstream homogeneous refinement. One potential solution is to jointly estimate the poses and the conformations^12^; however, a rigorous validation scheme is needed to exclude any reference bias in the process. Third, cryoSTAR correlates the coarse-grained model with 2D projections using a Gaussian density (see Methods for details). This approximates an ideal density map with distortion, which limits the ability of cryoSTAR to model small movements at higher resolution. Lastly, as discussed by other works^11,14^, the field lacks a method to evaluate and validate the resolved motions. Although a ground truth is nearly impossible to obtain, an FSC-style metric that quantifies the self-consistency or the level of confidence would be very useful for the users, preventing the over-interpretation of the results.

To further advance the capabilities of cryoSTAR, several directions can be explored in future work. For example, the simplification of each residue in cryoSTAR to one Gaussian bead from the coarse-grained model presents an opportunity for extension to finer-grained models, albeit with necessary modifications. Moreover, following the structural regularization technique, it is also possible to introduce more detailed regularizations that correspond to certain molecular dynamic (MD) force fields^28^. Integrating these energy-based regularizations presents a promising direction for uncovering hidden dynamics, though this approach still needs more exploration.

## Acknowledgments

We thank Dr. Zaixiang Zheng, Dr. Yiqun Wang, and Dr. Dongyu Xue for their insightful discussions on the project. We would like to thank Dr. Hang Li for the feedback on the project that helped shape this study. We also thank Dr. Michael Cianfrocco for the suggestions on the manuscript.

## Code availability

CryoSTAR software is freely available at https://github.com/bytedance/cryostar under the Apache Li- cense, Version 2.0.

## Methods

The main goal of cryo-EM reconstruction is to reconstruct a 3D volume *V* ∈ ℝ^*D*×*D*×*D*^ from its 2D projections {*I*^(*i*)^ ∈ ℝ^*D*×*D*^|*i* = 1, 2, ⋯, *M*}, where *D* is the side length and *M* is the size of dataset. By associating an atomic structure with a volume, cryoSTAR outputs a coarse-grained atomic structure *S* ∈ ℝ^*N*×*3*^ where *N* is the number of residues. Figure 1 shows the outline of cryoSTAR, which can be divided into five parts: (i) Given an input particle image *I*^(*i*)^ ∈ ℝ^*D*×*D*^ (the superscript *i* will be omitted for the simplicity of the notations), a variational autoencoder (VAE)-based deformation prediction network first infers the CG structure *S* ∈ ℝ^*N*×3^ of a molecule; (ii) We then convert the backbone structure *S* to an volumetric Gaussian density *V* ∈ ℝ^*D*×*D*×*D*^ and calculate its 2D projection ^*I*^^ ∈ ℝ^*D*×*D*^; (iii) The output backbone model is constrained by structure-aware regularization; (iv) We combine it with the auto-encoding supervision between *I* and ^*I*^^ to form the overall loss function; (v) The learned latent variable *z* is detached from the VAE and used to train a volume decoder to obtain a neural network representation of the density maps *V*’ ∈ _ℝ_*D*×*D*×*D*.

### Predicting the deformations with a VAE

CryoSTAR first uses the coordinates of the Cα of the amino acid or the P of the nucleotide as the center of the corresponding residue to build a coarse-grained model *S*_ref_ ∈ ℝ^*N*×3^ from a given a reference atomic model, where *N* denotes the number of the residues. Given an image *I* ∈ ℝ^*D*×*D*^ from the cryo-EM dataset, cryoSTAR uses a VAE to predict a corresponding deformation Δ^*V*^^ that modifies the reference structure to the deformed structure ^*V*^^ = *V*_ref_ + Δ^*V*^^. The encoder ℰ: ℝ^*D*×*D*^ → ℝ^|*z*|^ and decoder *D*: ℝ^|*z*|^ → ℝ^*N*×3^ are both multilayer perceptrons (MLPs). MLP is a global feature extractor and we do not observe any performance gain of CNN in practice. The hidden dimensions of the encoder and decoder are set to (512, 256, 128, 64, 32) and (32, 64, 128, 256, 512) respectively, and the latent space has 8 dimensions.

### Correlating density maps to atomic models with Gaussian densities

We use *P*: ℝ^*N*×3^ → ℝ^*D*×*D*^ to represent a physics-aware projection module which computes the projection from a given orientation. Specifically, the projection module *P* first converts the deformed molecular structure ^*V*^^ ∈ ℝ^*N*×3^ into a volumetric representation ^*S*^^ ∈ ℝ^*D*×*D*×*D*^, and then computes the projections ^*I*^^ ∈ ℝ^*D*×*D*^ with a given particle pose. Specifically, cryoSTAR uses a combination of Gaussian functions to correlate a volumetric density map to a coarse-grained atomic model. The volumetric density can be defined as the summation of *N* Gaussian pseudo-atoms, each denoting a residue’s electronic density in the reference atomic model:

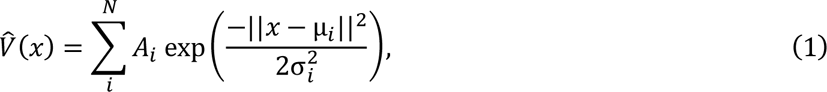

where *x* ∈ ℝ^3^is a coordinate in the sampling space, *μ*_*i*_is the *i*-th residue in the predicted structure ^*V*^^, *A*_*i*_ and σ_*i*_ are the amplitude and the variance of the Gaussian pseudo-atom. In detail, the amplitude *A*_*i*_is determined based on the total electrons of the corresponding amino acid or nucleotide, and the width σ_*i*_ is set to 2 empirically. The conformational heterogeneity is thus modeled by the relocation of the Gaussian pseudo-atoms.

Obviously, the Gaussian density is not equivalent to the density map reconstructed from the particles. To mitigate this distortion, we calculate the Fourier shell correlation (FSC) between the Gaussian density and the consensus density map, and low pass filter the Gaussian density to the frequency where *FSC* = 0.5, unless otherwise noted. In other words, cryoSTAR only uses low to intermediate resolution information to help resolve the continuous heterogeneity in cryo-EM data. This is also the reason that a more fine-grained model (e.g., a full-atomic model) is not used in this study. Given a volumetric density ^*S*^^, the projector further projects it to a 2D image ^*I*^^ by the image formation model:

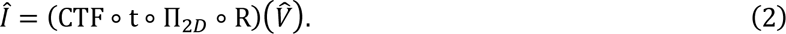

Here, R ∈ *SO*(3) is the orientation angle, Π_2*D*_ is a projection operator, t is the in-plane translation, and CTF is the contrast transfer function (CTF). These parameters can be obtained from previous processing steps and are assumed to be known in this study. Therefore, the projector is *differentiable* but *non-trainable*.

### Structural regularization

For the sake of brevity, we only consider biomolecules with a single, continuous chain with length *N* in our formulation and use subscript *i*, *j* to index its residues. We use *d*_*ij*_ and ^*d*^^_*ij*_ to denote the distance between the *i*-th and *j*-th residues in the reference structure *S*_ref_ and the predicted structure ^*V*^^, respectively. We keep this convention for all structural regularization definitions. It is worth noting that the structural regularization can be applied to more complex cases such as structures with multiple chains, missing residues, etc., as shown in Results.

### Incorporate basic structure information

The sequence of the target molecule should be unchanged in any conformational dynamics. The continuous loss ℒ_cont_ enforces the connection between two adjacent residues to be intact, which is defined as:

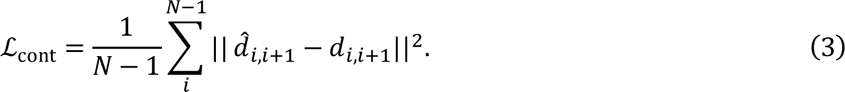

This term enforces the two adjacent residues to maintain their connectivity. To prevent two residues from clashing after predicting the deformation, cryoSTAR calculates a clash loss ℒ_clash_ on the pairs of residues that ensure the following condition during training:

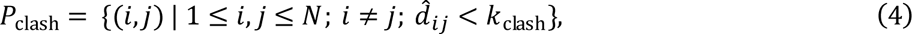

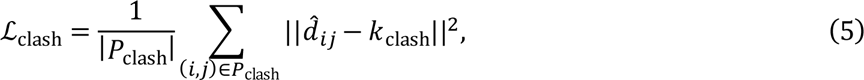

where *P*_clash_ denotes the set of residue pairs that experience collision during training. (*i*, *j*) is an index pair numbering the residues in the structure. Results will be penalized when the two non-adjacent residues are closer than the length *k*_clash_, preventing residue clashing from happening. We set *d*_clash_ to 4 Å in this paper. For a multi-chain complex, we consider all the residues including those from other chains for the calculation of ℒ_clash_.

### Preserving local rigidity with elastic network

CryoSTAR encourages the local structure to remain rigid. For example, an alpha-helix should keep its shape without being sheared, stretched, or destroyed in conformational dynamics. This is reasonable because: (i) modification of the local structure (secondary structure) is rare in the conformational dynamics captured by cryo-EM; (ii) when changes in secondary structure do occur, they are typically resolved through discrete 3D classification methods. CryoSTAR parameterizes the preservation of the local shape and geometry of the backbone model with an elastic network (EN). Specifically, given the initial structure, cryoSTAR builds an elastic network by connecting the residue pairs within a pre-defined distance. The elastic network loss is defined as:

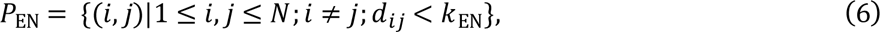

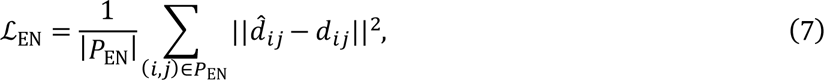

where *P*_EN_ is a set of edges for building the elastic network, and the constant value *k*_EN_is a predetermined cutoff. The goal of the elastic network is to preserve the local structural information in the reference atomic model. The connection between two Gaussian pseudo-atoms implies the possible non-covalent interactions between the two residues. In this paper, we set *k*_EN_ to 12 Å, which is a commonly chosen distance to model non-covalent interactions in a coarse-grained model^29^. For a multi-chain complex, we do not build edges between different chains.

### Sufficient conformation searching by adaptive relaxation

A static elastic network may be subject to the bias from the given reference atomic model, while conformational change often involves the forming and breaking of certain non-covalent interactions. For example, using the closed state of adenylate kinase (PDB: 1AKE) as the reference atomic model, the elastic network will cause the *“interlock”* problem, making it impossible to transition to the open state (PDB: 4AKE) (Extended Data Fig. 1).

To identify and mitigate these undesirable interactions, cryoSTAR adaptively selects the edges presented in the elastic network for regularization. To be specific, in each mini-batch of size *b*, cryoSTAR predicts *b* distance values, ^*d*^^_*ij*_’s, for each edge in the elastic network defined above, where (*i*, *j*) ∈ *P*_EN_. Subse-quently, we compute the variance of the predicted distances for each edge, denoted by Var(^*d*^^_*ij*_), over the set of these *b* values. We posit that the variance of the edge distance reveals the stability of edges during training, with higher variance indicating a greater likelihood of edge disruptions. All variances corresponding to the edges in *P*_EN_ can be grouped into a set *V* ≜ {Var(^*d*^^_*ij*_)|(*i*, *j*) ∈ *P*_EN_}. In the calculation of the loss, we only retain edges with variance below a certain percentile threshold *p*. Thus, only a subset of *P*_EN_ is used for the calculation of loss ℒ_EN_:

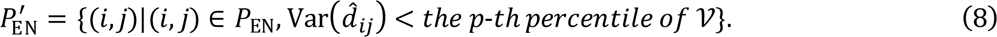

Note that *P*^′^ changes for every iteration. Unless otherwise noted, *p* = 0.99 in this paper. The rationale behind this is that the interactions responsible for stabilizing secondary structures should be unchanged, while the interactions involved in the conformational changes are unstable, exhibiting greater variance during training. Eliminating these unwanted constraints encourages the model to explore a wider range of potential conformations supported by the data. This allows cryoSTAR to mitigate the potential bias stemming from the reference atomic model.

### Loss function

The final loss function ℒ is composed of three parts: (i) the image reconstruction loss ℒ_image_, (ii) the structural regularization loss ℒ_struct_, and (iii) a posterior regularization term ℒ_KL_, which encourages the latent code *z* to be normally distributed.

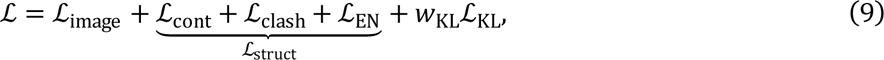

where 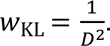

For the image reconstruction loss, cryoSTAR uses cross correlation to measure the similarity between an input image ^*I*^^ and the predicted projection ^*I*^^:

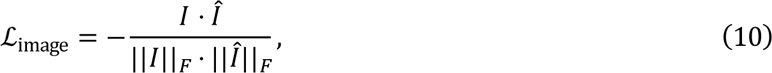

where ⋅ denotes a pixel-wise multiplication; ||*I*||_*F*_ is the Frobenius norm of *I*, which is a constant number and can be omitted.

### Generating density maps with the volume decoder

After the VAE is fully trained, the latent variable is detached and used to train a volume decoder to obtain a neural network representation of the density maps. The volume decoder models a volume in the Hartley space (similar to the Fourier space), following cryoDRGN^11^. It utilizes a multi-layer perceptron (MLP) to map a latent variable (representing a conformation) and a 3D coordinate to a value. The coordinate is encoded with positional embedding composed of 3*n* sinusoids (*n* is set to the side length of the particle) whose frequency is Gaussian distributed^30^. In the training phase, for each particle and the corresponding latent variable, we extract a slice (determined by the given pose) of the volume and compute the mean-squared-error (MSE) in the Hartley space. In the inference phase, we evaluate the total volume and transform it into the real space to obtain the density map.

### Latent space sampling

The auto-encoding framework enables us to obtain the latent variables describing the heterogeneity. Particles with similar conformational states will gather and stay away from other states during training. To visualize the significant conformational changes present in the data, we perform PCA on the posterior latent space and sample along the principal components axes to display the generated results. Specifically, in this paper, we uniformly sample 4 (shown in the Figures) or 10 points (shown in the movies) from the 5th to 95th percentiles along the principal component axes. It should be noted that we take the point in latent space closest to the point on the principal component axes to ensure that the sampling result is supported by data.

### Validation on the synthetic dataset

We validate cryoSTAR on a synthetic dataset with known ground truth. We use *E. coli* adenylate kinase (AK) as the target protein; it is known that it has both open and closed states with existing atomic models (pdb: 4AKE and 1AKE). We generate a 50-frame trajectory between the atomic models of the closed (pdb: 1AKE) and open (pdb: 4AKE) states using PyMOL, use the *e2pdb2mrc* program in EMAN2^31^ to compute the density maps from the full-atomic models, and generate synthetic particles with the statistics in Table 1 (Extended Data Fig. 1a). The closed state model (pdb: 1AKE) is used as the reference structure. Since the protein is much smaller than the experimental dataset, *p* for the adaptive relaxation is set to 0.8.

**Table 1.** Statistics of the synthetic dataset.

| Parameters | Value |
| --- | --- |
| Number of particles | 50,000 |
| Image size | 128 px × 128 px |
| Image pixel size | 1.0 Å |
| Defocus | Lognormal(1.0, 0.3<sup>2</sup>) μm |
| Accelerating voltage | 300 keV |
| Spherical aberration | 2.7 nm |
| Amplitude contrast | 0.1 |
| SNR (with Gaussian noise) | 0.0001 |
| Pose/rotation | Uniform on SO(3) |
| Pose/translation | 0 |

By traversing across the learned latent space, we can generate the corresponding density maps and the backbone models. CryoSTAR correctly identified the motion of AK between the open and closed states (Extended Data Fig. 1b, 1e and 1f). Both generated density maps and coarse-grained models recovered the correct motion. The Cα-RMSDs between the generated models and the ground truth atomic models are around 2 Å (Extended Data Fig. 1c) across all the conformations, and the FSCs between the density maps and the ground truth densities are around 6 Å at the 0.5 cut-off (Extended Data Fig. 1d).

### Training details on the experimental datasets

Details of the experiment, including running time on 4 x NVIDIA Tesla V100 GPUs, can be found in Table 2. For all datasets, we first calculate the Fourier shell correlation (FSC) between the Gaussian density and the consensus density map, and low pass filter the Gaussian density to the frequency where *FSC* = 0.5 during Phase I training. The only exception is the TRPV1 channel dataset (EMPIAR-10059), since the FSC curve drops drastically at very low resolution due to the presence of the lipid nanodisc, therefore we manually set the low pass filter cutoff to be 10 Å. For the tri-snRNP spliceosome dataset (EMPIAR- 10073) and the pre-catalytic spliceosome dataset (EMPIAR-10180), particles are resized with Fourier cropping. For the α-latrocrustatoxin dataset (EMPIAR-10827), we first run a 2D classification on the particles in the given dataset and select the class averages showing only monomers.

**Table 2.** Training details and running times on the four experimental datasets.

| Dataset | Reference | # of residue | # of particles | Particle size (Phase I) | Particle size (Phase II) | Running time |
| --- | --- | --- | --- | --- | --- | --- |
| EMPIAR-10073 | 5GAN<sup>33</sup> | 9,325 | 138,899 | 128 px × 128 px | 256 px × 256 px | 5h4m |
| EMPIAR-10180 | 5NRL<sup>23</sup> | 13,941 | 327,490 | 128 px × 128 px | 256 px × 256 px | 7h54m |
| EMPIAR-10059 | 7RQW<sup>27</sup> | 2,128 | 218,787 | 192 px × 192 px | 192 px × 192 px | 18h30m |
| EMPIAR-10827 | 7PTX<sup>26</sup> | 1,000 | 230,218 | 224 px × 224 px | 224 px × 224 px | 3h6m |

### Comparison with other methods

One unique feature of cryoSTAR is its ability to generate both coarse-grained models and the corresponding density maps. As noted in the introduction, there are a few other methods that also use structural information from the atomic model and/or output backbone models. Namely, Zernike3D^19^ decomposes heterogeneity into a few bases with Zernike polynomials, and DeepHEMNMA^17^ uses NMA to maintain the structural integrity. NMA has also been used by a few other exploratory methods that are only applied to synthetic data^18,32^. However, these methods impose too strong regularization that prevents the model from exploring large motions. For example, the motions found by Zernike3D on the pre-catalytic spliceosome dataset (EMPIAR-10180) were significantly smaller than the results using other methods^19^. Similar results were also observed with NMA. In comparison, cryoSTAR only applies minimum and necessary regularizations. The adaptive relaxation further ensures that the model will not be severely biased by reference atomic model by “unlocking” some of the interactions.

CryoSTAR and cryoDRGN^11^ both use VAE as the base architecture. In cryoDRGN, the latent variable encoding heterogeneity is directly optimized from the data without any further regularization. This enables cryoDRGN to also resolve the compositional heterogeneity in the cryo-EM datasets. However, as cryoDRGN does not directly model heterogeneity as a deformation process, the trajectories generated often have artifacts where the flexible densities will vanish and then reappear (Extended Data Fig. 5a). Additionally, cryoDRGN struggles when dealing with membrane proteins unless preprocessed with particle subtraction, since it will focus on the regions with the highest variabilities, which usually are detergent micelle or lipid nanodisc (Extended Data Fig. 4a). Instead, cryoSTAR models conformational heterogeneity as the deformations of a reference model and ensures output integrity with proper structural regularization. The volume decoder of cryoSTAR is equivalent to the decoder of cryoDRGN, but the generated density maps are smoother with less artifacts in transition (Extended Data Fig. 5c). Moreover, cryoSTAR can solve conformational heterogeneity of membrane proteins without external masks or particle subtraction (Extended Data Fig. 4b), which is a big advantage compared to other methods.

3DFlex^14^ models conformational heterogeneity as an explicit deformation of a canonical density. As discussed in their paper, regularization plays a crucial role in 3DFlex, otherwise the results will suffer from various artifacts. 3DFlex applies a meshing scheme and defines a local rigidity regularization on the tetrahedral meshes to avoid overfitting on noise. In practice, an optimal mesh segmentation is important to ensure the best result. Moreover, since the optimization of the canonical density is strongly influenced by the deformation field found in the previous step, any overfitting upstream will lead to artifacts in the canonical density, and this cannot be detected with the “gold standard” FSC (Extended Data Fig. 5b). CryoSTAR echoes the idea of keeping local rigidity, but we explicitly define it on a coarse-grained model instead of on a volumetric density. Therefore, cryoSTAR minimizes the heuristics, and the structural regularization terms in cryoSTAR have more biological meanings. On the other hand, the volume decoder in cryoSTAR does not use any explicit deformation field as the input, so that the generated density maps are directly optimized from the particle images. In cryoSTAR, the trajectory of density maps while traversing in the latent space is not a series of convection of a canonical density. Last but not least, cryoSTAR is significantly faster than 3DFlex.

**Supplementary Video 1. Video of cryoSTAR results for the pre-catalytic spliceosome (EMPIAR- 10180).**

**Supplementary Video 2. Video of cryoSTAR results for the U4/U6.U5 tri-snRNP (EMPIAR-10073). Supplementary Video 3. Video of cryoSTAR results for the TRPV1 channel (EMPIAR-10059).**

**Supplementary Video 4. Video of cryoSTAR results for the α-latrocrustatoxin (EMPIAR-10827).**

## Supplementary Information

**Extended Data Figure 1.**
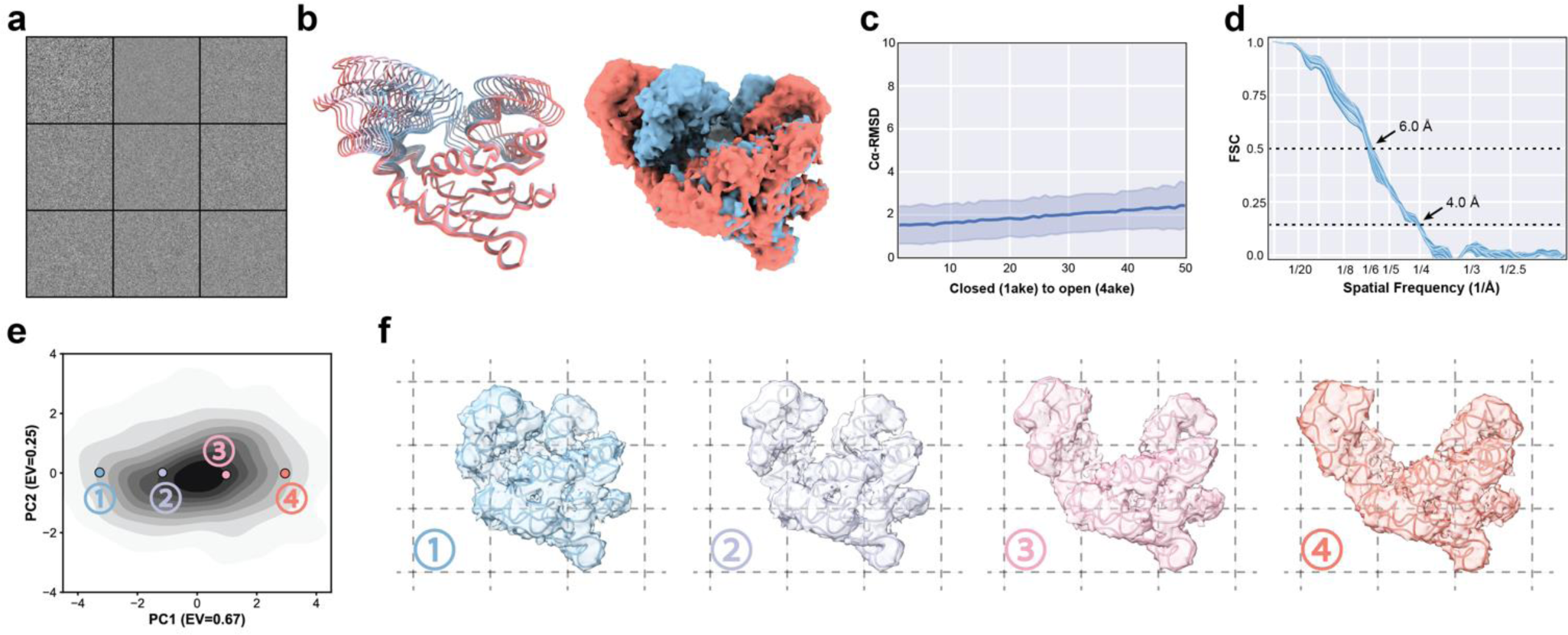
Validating cryoSTAR on a synthetic dataset (adenylate kinase), **a,** Examples of the particles from the synthetic dataset. A total of 50,000 particles were simulated from 50 continuous conformational states from closed to open (5,000 particles for each conformation). The signal-to-noise ratio is 0.0001. **b,** Colored series of coarse-grained models and particle density maps generated by cryoSTAR, sampling along the first principal component of the latent space, **c,** The Cu-RMSD between the output of cryoSTAR and the ground truth at different conformational states (left: closed state; right: open state). The middle curve denotes the mean and the light blue regions denote one sigma deviation, **d,** The FSC curves between the particle density maps and the ground truth densities at each conformation, **e,** PCA visualization of the cryoSTAR latent space, where the color depth represents the particle population, **f,** The coarse-grained models and particle density maps generated by sampling along the first principal component in the latent space, as marked in **e** with the corresponding colors.

**Extended Data Figure 2.**
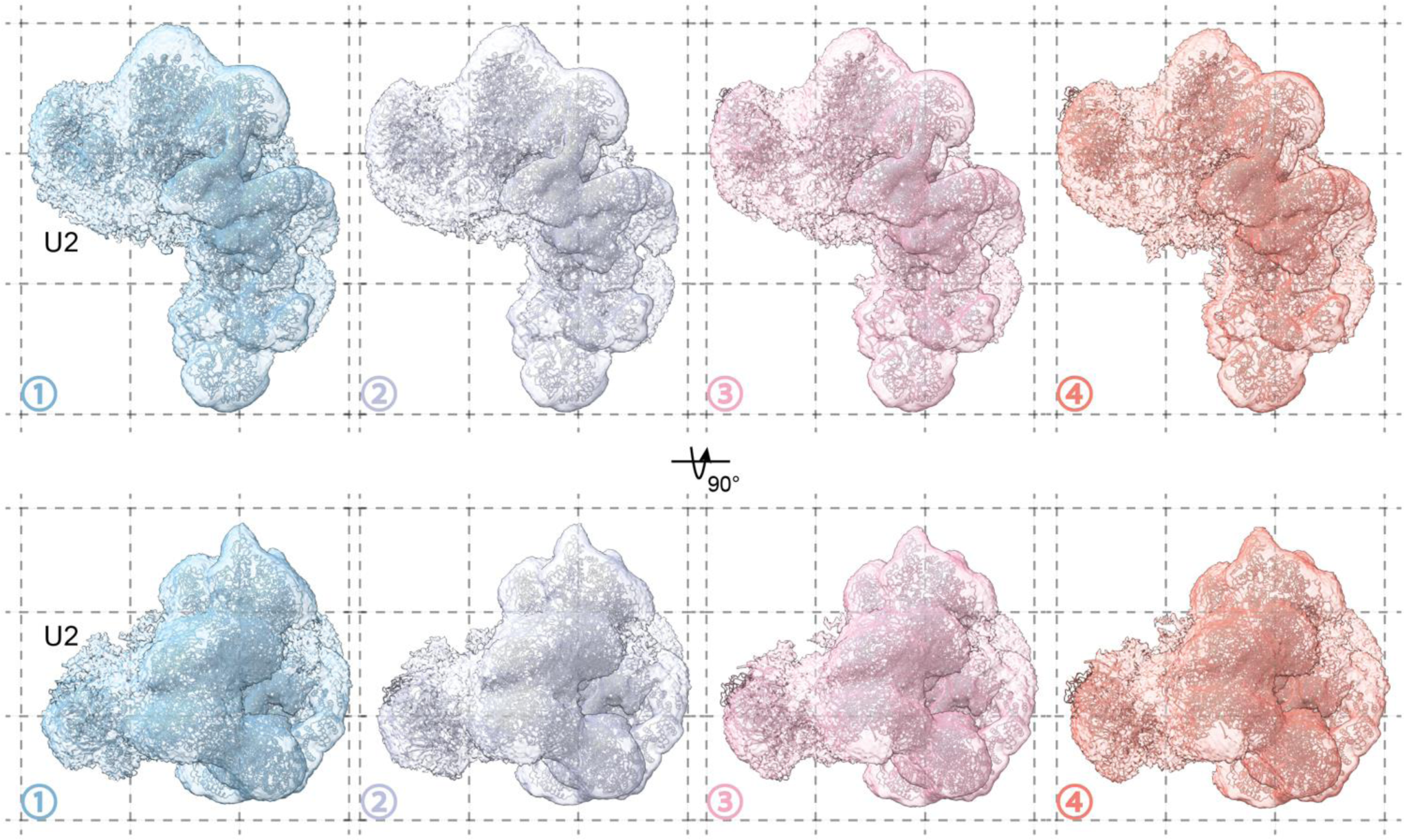
CryoSTAR applies to the pre-catalytic spliceosome dataset (EMPIAR-10180), showing the U2 core with a lower isosurface level. The coarse-grained models and particle density maps (top and side views) generated by sampling along the first PC in the latent space, as marked in Fig. 2b with the corresponding colors. The U2 core exhibits a weaker density compared to the densities elsewhere. Although the generated coarse-grained models implies a movement of the U2 core, this cannot be fully demonstrated by the generated particle density maps. All density maps are shown using the same isosurface levels, lower than the ones used in Fig. 2.

**Extended Data Figure 3.**
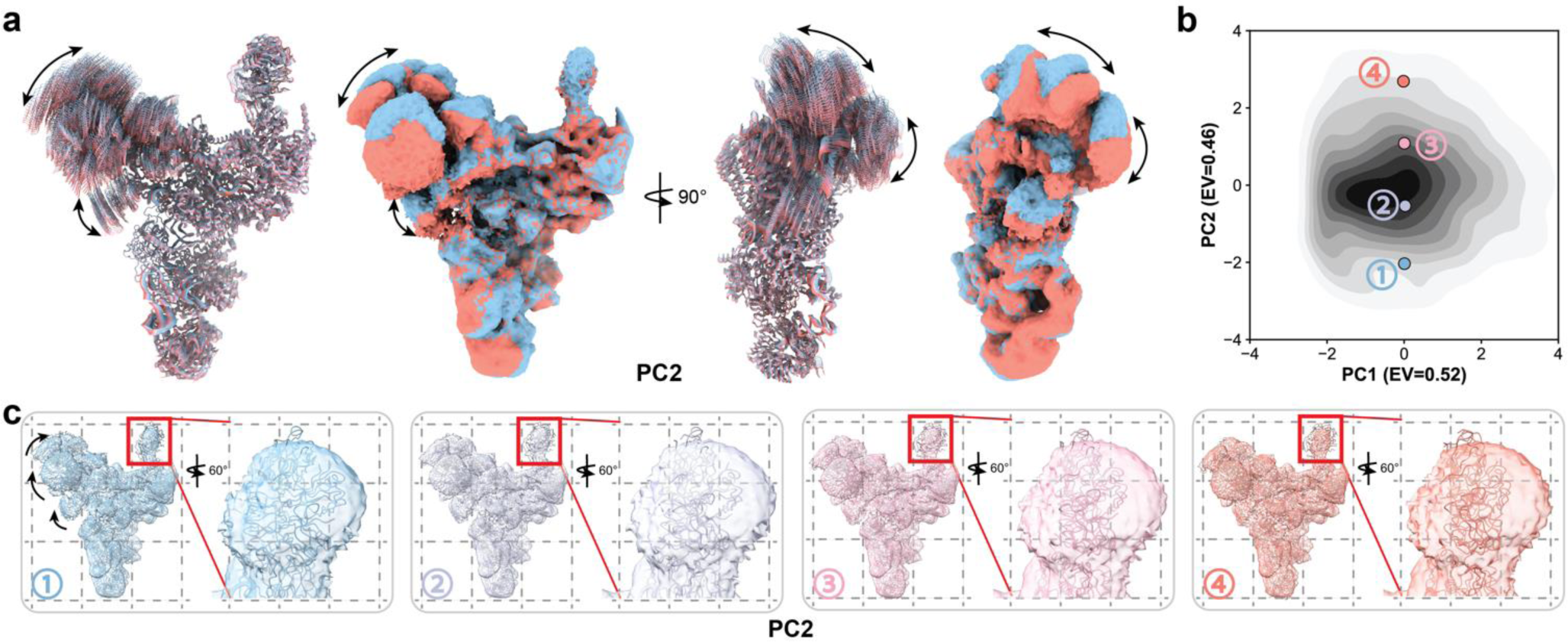
CryoSTAR applies to the U4/U6.U5 tri-snRNP dataset (EMPIAR-10073), showing the results traversing along the second principal component, **a,** Colored series of ten coarse-grained models and two particle density maps generated by cryoSTAR, sampling along the second principal component of the latent space, **b,** PCA visualization of the cryoSTAR latent space, where the color depth represents the particle population, **c,** The coarse-grained models and particle density maps generated by sampling along the second principal component in the latent space, as marked in **b** with the corresponding colors, and the side view of the structures zoomed in the arm domain, with a lower isosurface level. Although the generated coarse-grained models implies a rotation of the arm region, this cannot be supported by the generated particle density maps.

**Extended Data Figure 4.**
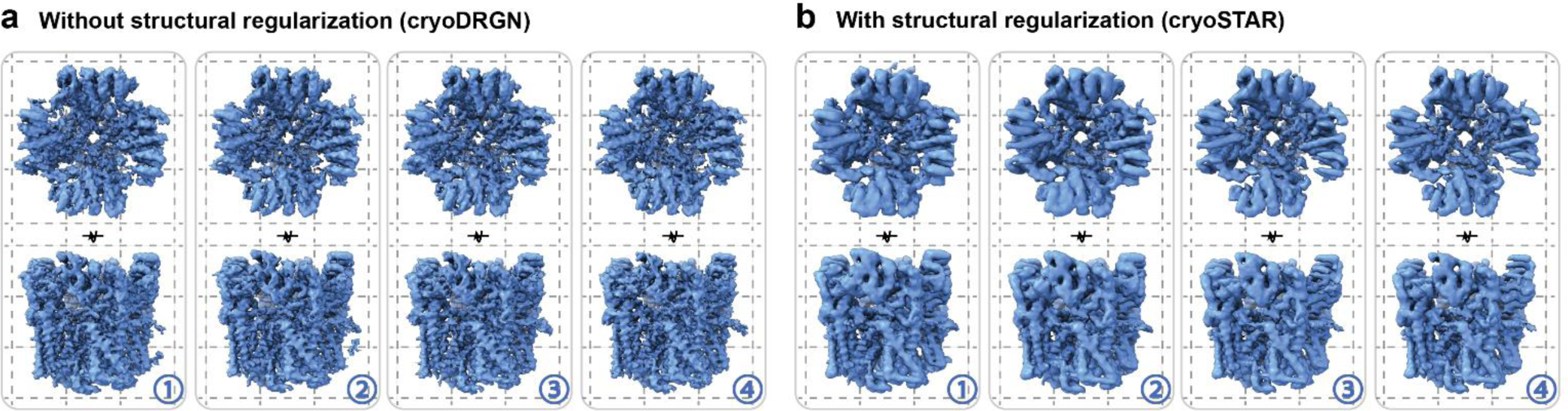
Structural regularization helps to reveal the dynamics in the TRPV1 dataset (EMPIAR-10059). **a,** Four density maps (top and side views) generated by cryoDRGN, a method without applying structural regularization, sampling along the first principal component of the latent space, using the same isosurface levels. CryoDRGN failed to reveal the motions in the TRPV1 dataset, because it focused on the area with the highest variability, which is the nanodisc region. This is typical when applying cryoDRGN on membrane proteins, **b,** With the help of structural regularization, cryoSTAR successfully uncovers the hidden motion in the TRPV1 dataset.

**Extended Data Figure 5.**
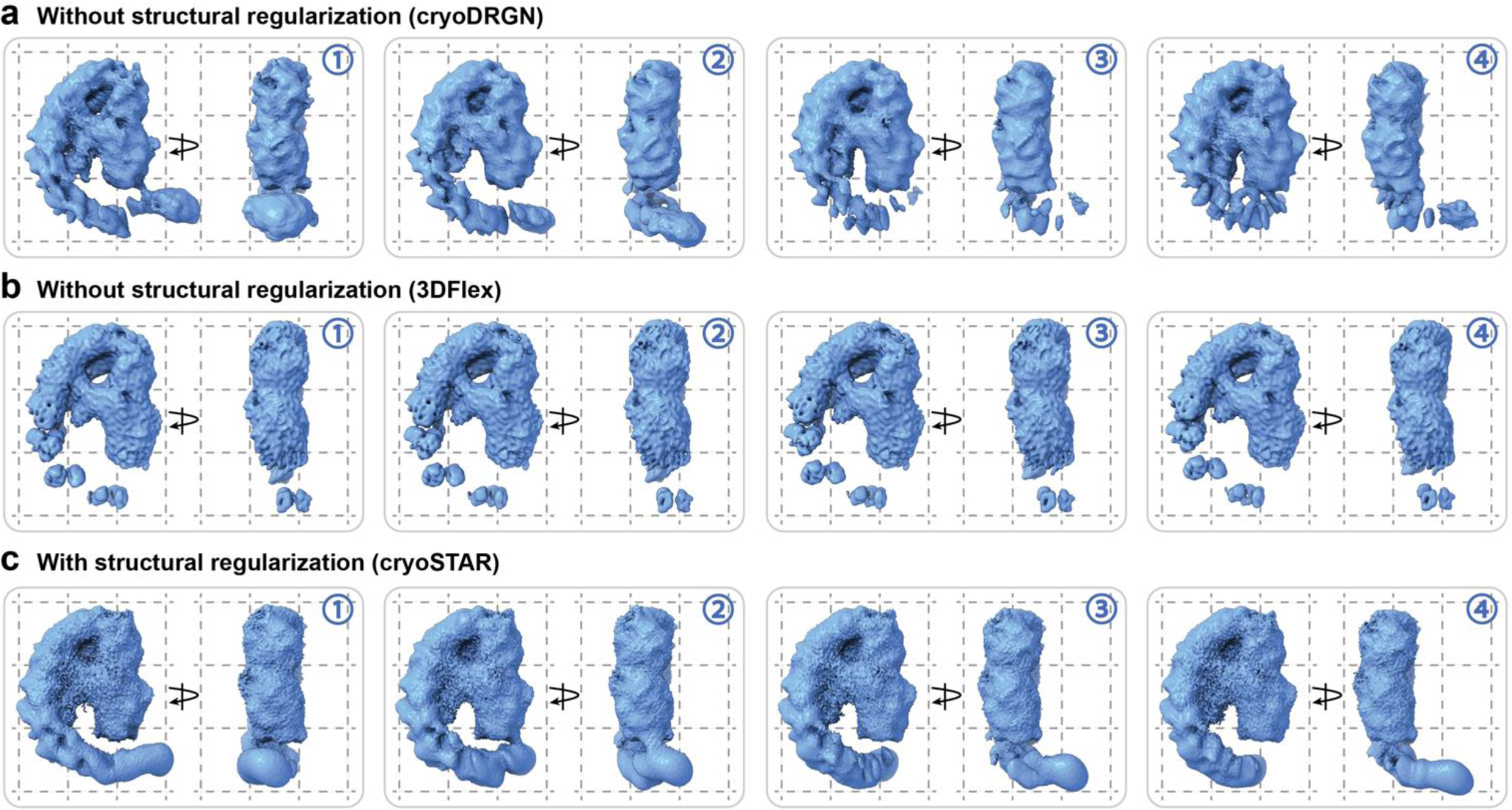
Structural regularization suppress the artifacts in the densities from the a-LCT dataset (EMPIAR-10827). **a,** Four density maps generated by cryoDRGN, sampling along the first principal component of the latent space, using the same isosurface levels. Without the structural regularization from the reference atomic model, the resulting output density may exhibit discontinuities and artifacts, including density disappearance while traversing along the latent space (3 and 4). **b,** Flex volume sampled along the first latent dimension from 3DFIex (default parameters). The estimated motion is much smaller. Despite the reported FSC resolution of 2.85 A, the flex densities are not continuous, with a severe artifact at the Ankyrin-like repeat domain (ARD), **c,** Structural regularization helps cryoSTAR to uncover the continuous motion without discernible artifacts in the reconstructed density maps.

## Footnotes

1 The short version of this paper has been accepted by the NeurIPS workshop on New Frontiers of AI for Drug Discovery and Development.

